# Mechanobiological cues to bone cells during early metastasis drive later osteolysis: a computational mechanoregulation framework prediction

**DOI:** 10.1101/2024.10.03.616269

**Authors:** Anneke S.K. Verbruggen, Elan C. McCarthy, Roisin Dwyer, Laoise M. McNamara

## Abstract

Bone cells contribute to tumour metastasis by producing biochemical factors that stimulate tumour cell homing and proliferation, but also by resorbing bone matrix (osteolysis) that releases further stimulatory factors for tumour growth in a vicious cycle. Changes in the local mechanical environment of bone tissue occur during early metastasis, which might activate mechanobiological responses by resident bone cells (osteocytes) to activate resorption (osteoclasts) and thereby contribute to tumour invasion. The objective of this study is to investigate whether bone osteolysis is driven by early changes in the bone mechanical environment during metastasis by (a) implementing subject-specific FE models of metastatic femora to predict the mechanical environment within bone tissue during early metastasis (3-weeks after tumour inoculation) and then (b) applying mechanoregulation theory to predict bone tissue remodelling as a function of the evolving mechanical environment within bone tissue during breast cancer-bone metastasis. We implemented a global resorption rate derived from an experimental model, but the mechanoregulation algorithm predicted localised bone loss in the greater trochanter region, the same region where osteolysis was prevalent after three weeks of metastasis development in the animal model. Moreover, the mechanical environment evolved in a similar manner to that reported in separate subject-specific finite element models of these same animals by 6 weeks. Thus, we propose that early changes in the physical environment of bone tissue during metastasis may elicit mechanobiological cues for bone cells and activate later osteolytic bone destruction.

## 1. Introduction

Breast cancer commonly migrates from the primary tumour and metastasises to bone, occurs for 70-80% of patients with advanced disease. It has been proposed that breast cancer cells have a particular affinity to the composition or physical properties presented by bone tissue, which provides an attractive environment for metastasis development according to ‘Seed and Soil’ theory [1]. Biochemically, bone cells play an important role in tumour metastasis. Osteoblasts express chemokines (CXCL12/SDF-1) that facilitate homing of tumour cells to bone. Tumour cells produce factors (PtHRP, MMP-1, IL-11, CTGF) that stimulate growth factor production by osteoblasts (e.g. TGF-β , osteopontin, RANKL), which induce tumour cell proliferation and also activate osteoclasts. Osteoclast resorption of bone matrix providing space for tumour invasion and releases growth factors (TGF β) from the matrix, which perpetuates osteolysis and tumour invasion. Thus, the biochemical interaction between cancer cells, osteoblasts and osteoclasts has been referred to as a ’vicious cycle’ [2].

The changes that arise in the bone microenvironment during bone metastasis are not limited to biochemical changes, because the composition, mechanical properties and local mechanical environment of the bone tissue are also altered [3–7]. Specifically, bone mineral content and cortical thickness were significantly altered by 3 weeks of tumour development and occurred prior to bone loss and the development of overt osteolytic lesions, which were not reported until 6 weeks [6]. In a following study, micro-CT-derived finite element (FE) models of these metastatic proximal femurs were applied to study the mechanical environment within bone tissue during bone metastasis [7]. Interestingly, in early metastasis there was a decrease in strain distribution within the bone tissue of the proximal femur, which coincided with the onset of cortical thickening and mineralisation of bone [6].

Osteocyte cells regulate bone resorption or formation by osteoblasts and osteoclasts depending on the mechanical demands on the tissue. The biochemical growth factors involved in mechanically activated bone cell signalling (PTHrP, RANKL, TGF-β, Ca ) are also those that attract invading cancer cells and influence resident osteoclasts to activate bone resorption and osteolysis. Thus, mechanobiological signalling by osteocyte cells, initiated because of the altered mechanical environment, might contribute to the vicious cycle by activating osteoclast resorption, and thereby play a role in the cancer vicious cycle. However, the role of bone cell mechanobiological responses in metastatic osteolysis is not yet understood.

To date, studies have largely sought to understand the biochemical factors by which bone cells influence cancer proliferation and the viscous cycle that ensues when the cancer cells then produce regulatory factors (PTHrP, RANKL) that further drive osteoclastic bone resorption. However, how bone remodelling is affected by the evolving mechanical environment that the cancer cells initiate is not well understood. Mathematical and computational models have been applied to simulate and predict the role of biochemical processes and cell-cell signalling in tumour growth, and to a lesser extent have considered biophysical processes [8–13]. Bone remodelling has been studied by the development of a number of mechanoregulation theories, which predict bone remodelling and adaptation on the basis of mechanical stimuli such as strain, microdamage and fluid velocity [14, 15]. Osteocytes experience fluid shear stress within the lacunar-canalicular network under compressive loading during normal physiological movement, and also extend protein attachments to their matrix and thus can experience matrix strain (tensile or compressive). Mechanoregulatory responses are driven by a mechanical stimulus, and previous studies have assumed that stimulus to be the strain energy density tensor [14, 16] to account for the evolving Young’s modulus, density and strain. Such approaches have been applied to study bone resorption and formation of healthy and diseased bone tissue in combination with micro-CT derived finite element models [17–24]. However, mechanoregulation theory has not yet been applied to investigate the contribution of changes in the mechanical environment for driving bone resorption during the cancer vicious cycle. Applying such approaches may shed light on the potential role of mechanobiology in the development of osteolysis as bone metastasis progresses.

The objective of this study was to test the hypothesis that an altered physical environment of bone tissue during early metastasis could elicit mechanobiological cues for resident bone cells and thereby contribute to later osteolytic destruction. We implemented (a) subject-specific FE models to predict the mechanical environment within the bone tissue at the early stage of metastasis and (b) a mechanoregulation theory based on strain energy density to predict how material properties are altered as a function of the evolving mechanical environment within bone tissue during breast cancer-bone metastasis.

## 2. Methods

In this study an iterative approach is taken, whereby subject-specific FE models are implemented to predict the mechanical environment within the bone tissue at the early stage of metastasis. A mechanoregulation approach, which adapts the bone tissue properties on the basis of the mechanical environment (strain energy density), is applied to predict how material properties are altered as a function of the evolving mechanical environment within the bone tissue.

### 2.1. Finite element model development

The current study builds upon µCT-derived FE models, developed in a previous computational study of strain distribution during breast cancer metastasis [7]. In brief, immune competent BALB/c mice, inoculated into a femur-adjacent mammary fat pad with 4T1 breast cancer cells, were euthanised, at either 3 or 6 weeks post-inoculation, and µCT scanning and reconstruction was conducted to develop solid models of the proximal femora.

In the current study, finite element models of the 3-week murine cohort were developed by meshing the reconstructed solid models with 4-noded linear elastic, homogeneous, tetrahedral elements, following mesh convergence analysis, using 3Matic and Abaqus software (Figure 1, 1B). A study compared 4- and 10-noded tetrahedral meshes in a human proximal femur FE model under compression and reported no significant differences in their output stress, strain or overall accuracy compared to experimental results [25]. Loading was applied to the femoral head surface equating to 120% of bodyweight, recorded for each animal during the experimental study [6] and applied in the proximal-distal direction and 10.9% in the posterior-anterior direction to reflect the peak loading mid-way through a murine trotting cycle [26] (see Figure 1C). The distal surface was fixed in all directions (**Figure 1 Error! Reference source not found.** C). For the current study, each computational model was assumed an initially homogeneous, isotropic structure, with the mean bone mineral density assigned according to µCT imaging of the metastatic femoral bone (1.262 ± 0.02 g/cm^3^), converted to ash density [6] and the corresponding Young’s modulus applied according to a power law relationship used in female mouse tibiae, according to the following equation [27].

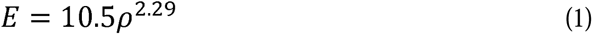

**Figure 1:**
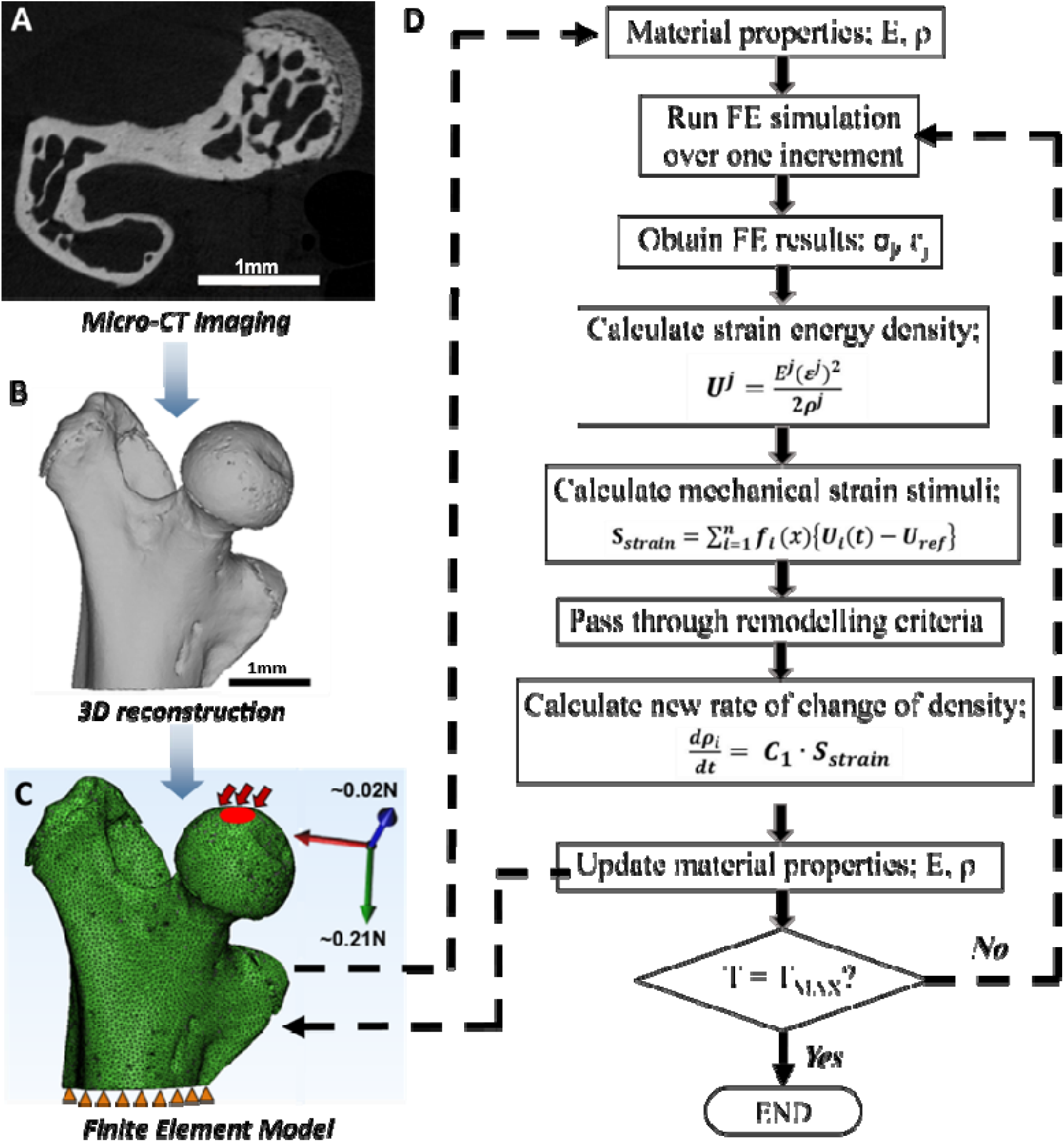
Flowchart outlining approach to combining micro-CT derived finite element modelling and mechanoregulation theory to predict bone remodelling in metastatic bone tissue. (A) µCT imaging was conducted on the proximal femur of metastatic mice at 3 weeks post-inoculation of breast cancer cells. (B) 3D reconstructed models were generated by segmenting µCT scans. (C) Loading and boundary conditions were applied to the femoral head surface (in red) and fixed distal surface (in orange) to reflect physiological loading in a mouse hindlimb. (D) The mechanoregulation algorithm was applied iteratively over a period of 21 days (T_MAX_) to predict bone remodelling on the basis of strain energy density.

### 2.2. Mechanoregulation Theory

To introduce mechanoregulation theory, a user-defined field subroutine was applied, whereby the density and Young’s modulus of each element within the model mesh was incrementally updated according to the rate of change of density predicted by a remodelling algorithm.

Specifically, density adaption was predicted on the assumption that a mechanoregulatory response is driven by a mechanical stimulus, strain energy density tensor, U ^j^ [14, 16], calculated according to:

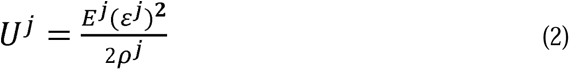

where, at location *j*, the tissue Young’s modulus, *E ^j^* (MPa) and density, ρ*^j^* (g/cm^3^) are defined and the strain (□*^j^*) is predicted by solving the finite element model. The stimulus, *S_strain_*, is calculated by comparing to the reference strain energy density, *U^ref^*, which is calculated at homeostatic equilibrium, according to the following equation:

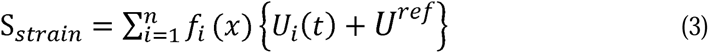

The homeostatic region was defined according to Frost’s ‘Mechanostat’ theory [28], adapted for bone tissue mechanoregulation. Reference strain energy density, *U^ref^*, is calculated as a function of reference modulus *E^ref^*, density *^ref^* and maximum principal strain *^ref^*, when bone remodelling is at equilibrium (see equation 4). This ‘lazy zone’ of equilibrium is defined by strain thresholds for net bone resorption and formation are □ and □ , respectively. The value of L is thus determined by the current value of □ , according to conditions outlined below:

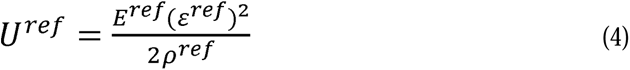

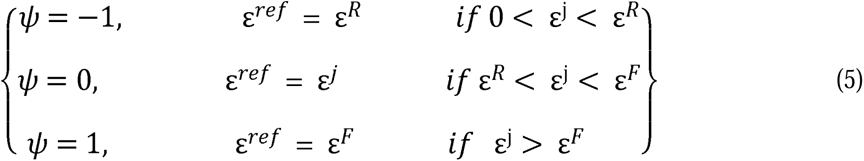

where ψ is the mechanical signal ensuring density decreases under resorption conditions and increases upon bone formation. Finally, the rate of change of density in response to mechanical stimuli, *d*ρ*i/dt*, and a new resulting density, (ρ*_i+1_)*, are calculated using the following equations:

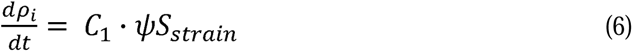

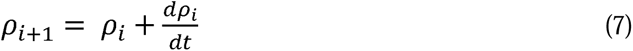

where a time constant, *C*_1_, governs the rate of remodelling. This constant was determined separately for bone tissue resorption (C_1_^R^ = 1600) and formation (C_1_^F^ = 1.79) as described below, and the rate applied was dependent on the strain threshold as follows:

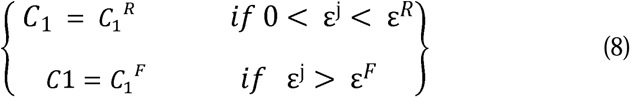

Maximal and minimal apparent density limits of 1.73 g/cm^3^ and 0.01 g/cm^3^ were assigned as standard, where density below 0.01 g/cm^3^ represents complete resorption [29, 30]. These equations (1 – 7) are thus implemented sequentially as part of the remodelling algorithm to iteratively predict a new density and Young’s modulus for each element within a given model mesh, as illustrated in a flowchart (Figure 1D).

### 2.3. Determining bone remodelling thresholds and constants

An upper strain threshold, beyond which bone tissue undergoes formation, is commonly reported to be approximately 1000µε [31–33] and was applied to the remodelling algorithm ε for this study. In contrast, a wide variety of resorption threshold values have been theorised, ranging from 27.9 - 250µ [21, 28, 34]. Considering this range of thresholds, we conducted a parameter variation study (Supplementary Fig 1B) to determine the lower strain threshold (100µε), below which bone resorption would occur.

We first derived the remodelling rates (C_1_) for the mechanoregulation theory by considering a simplified model of an individual bone trabecular strut under loading, such that the average change in bone mineral density (M_mean_) was representative of that measured experimentally for trabeculae of the proximal femur after 6 weeks of metastasis in the animal model [6]. Briefly, the trabecular strut model was assigned material properties obtained experimentally for trabecular metastatic bone after 6 weeks of metastasis [6] (E = 14.117 GPa, ν = 0.3, ρ = 1.138 g/cm ) and to minimise edge effects a surrounding ‘non-bone’ region (E = 0.001GPa, ν = 0.3, ρ = 0.001) was included, see Supplementary Figure 2A. The axial surface of the strut was fixed in all directions and a controlled low-strain (25μ ) displacement was applied to the opposing face in the longitudinal direction (SupplementaryFigure 2B). The time constant (C_1_) was derived to predict bone formation (C_1_^F^) undergoing an overall density increase (1.123 g/cm^3^ to 1.178 g/cm^3^) at high strain (1250 ) over 3 weeks (21 increments), which was based on density increase in healthy trabecular bone in the experimental study [6], see Supplementary Figure 2D. The rate for bone resorption (C_1_^R^) was derived such that an overall density decrease (1.138 g/cm^3^ to 1.128 g/cm^3^) would occur over the same period, which was based on experimental measurements of the average decrease in bone mineral density (M_mean_) during resorption of metastatic trabecular bone [6]. A parameter variation study was then performed for *C_1_^R^* (Figure ***2***), which revealed that a value of 1.6x10^3^ predicts bone resorption within the greater trochanter regions, similar to experimental observations of overt osteolytic degradation in the greater trochanter region, over the same time period, reported previously [6]. Higher *C_1_^R^* values inaccurately predicted the majority of the proximal femur model to be resorbed (Figure ***2***). Therefore, *C_1_^R^* = 1.6x10^3^ was applied for all further models.

**Figure 2:**
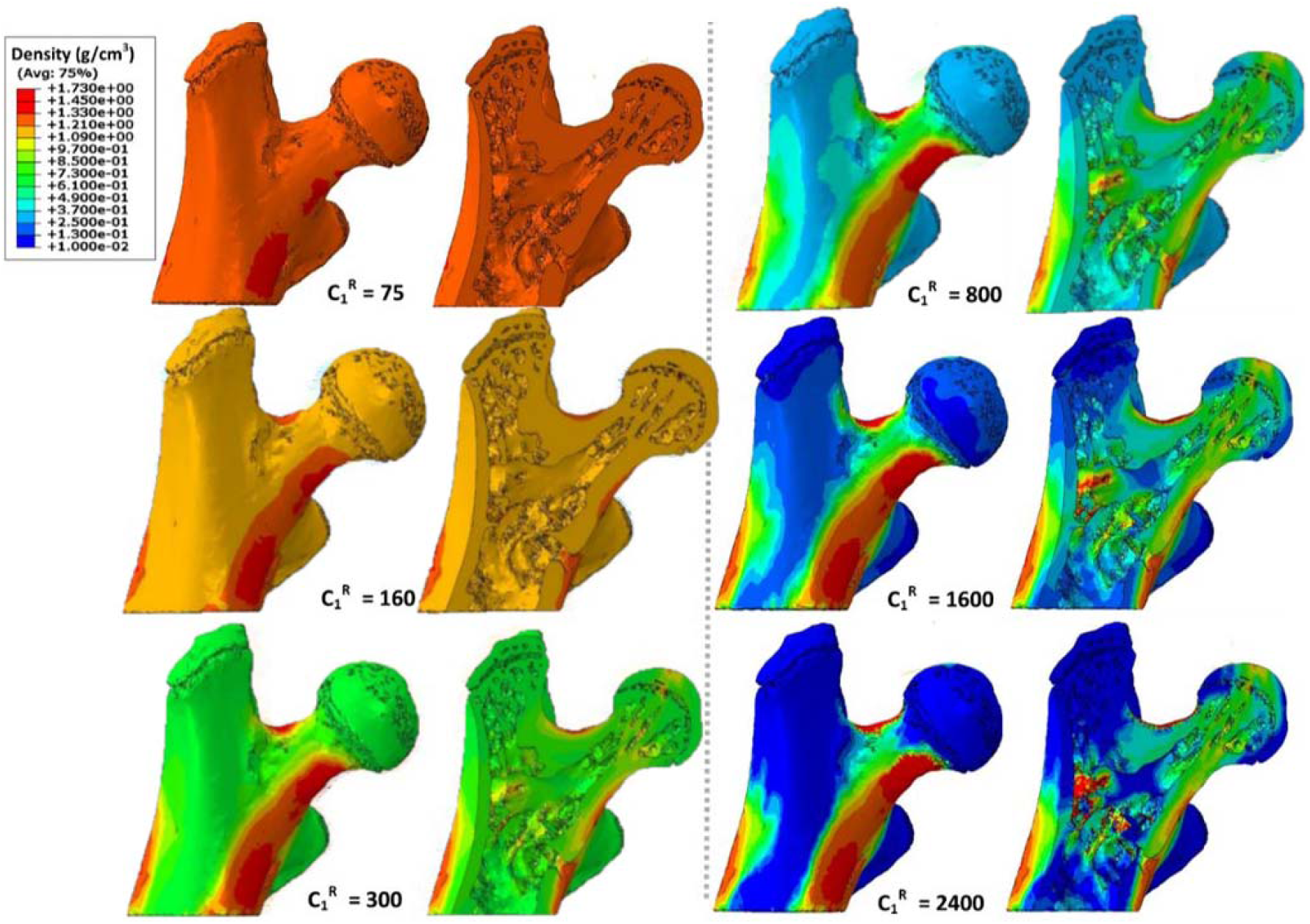
Bone resorption constant parameter variation study. Predicted bone mineral density distributions for a metastatic model (Whole bone – left, cross-section – right) after 3 weeks application of the mechanoregulaiton theory with different resorption constants (C_1_^R^) . A resorption rate of 1600 was deemed appropriate to represent the global rate of resorption observed in the experimental study that informed model development [6].

### 2.4. Finite Element Analysis

This study sought to predict osteolysis in response to changes in the mechanical environment of metastatic bone, using finite element analysis (FEA) and mechanoregulation theory, and compare these to findings from previous experimental and computational studies. Qualitative analysis of the distribution of bone tissue density and strain energy density (SED) throughout the proximal femur was performed. In terms of quantitative analysis, maximum principal strain distribution was assessed, along with maximum principal strain and SED in these models. Density resorption was also analysed quantitatively, by calculating the percentage volume of each model that had decreased in density throughout the remodelling process. The decrease in density was considered notable if resorbed below 0.1 g/cm^3^, an interim value between mean model density (1.262 ± 0.02 g/cm^3^) and complete resorption (0.01 g/cm^3^), whereby elements with a value below this threshold have reduced in density by over 90%.

Changes in bone tissue density and maximum principal strain within the micro-CT derived finite element models of metastatic (MET) proximal femurs at 3 weeks, and the predicted changes arising by 6 weeks after application of the mechanoregulation theory, were analysed. These results were compared to results of our previous FE study [7], in which micro-CT derived FE models from the 3 week timepoint were identical except for their heterogeneous distribution of density, and also included a separately imaged 6-week cohort of metastatic proximal femur heterogeneous models for direct comparison. The subscript, ***m***, describes results from homogeneous FE models with applied mechanoregulation theory (i.e. findings from the current study), while subscript ***s*** denotes the heterogeneous FE models imaged separately at 3 weeks and at 6 weeks post-inoculation [7]. In this way, MET_het_ models present the mechanical environment of bone tissue at two time points of the study and can serve as a direct comparison for the 3-week and 6-week time points predicted by the bone-remodelling algorithm, determined according to mechanoregulation theory (MET_hom_).

### 2.5. Validation of models

The initial homogenous models were validated against subject-specific heterogeneous finite element models, which were described previously (Verbruggen and McNamara 2023). Briefly, micro-CT (5μm) data for bone mineral density from these bones at the initial time point (3 weeks post-inoculation) were analysed to determine material parameters according to a power law relationship, which related gray value, mineral density and Young’s modulus. This approach provided 100 distinct uniformly distributed properties to capture the heterogeneity from the micro-CT data. The loading and boundary conditions were applied as described above. We compared the predicted strain and strain energy density of the homogeneous models (3wk METhom) to these subject-specific heterogeneous models (3wk METhet) for two different resorption rates (1600, 160), see Figure 3 and Supplementary Figure 3. The predictive ability of the mechanoregulation theory was also validated against published experimental data, see Supplementary Figure 4.

**Figure 3:**
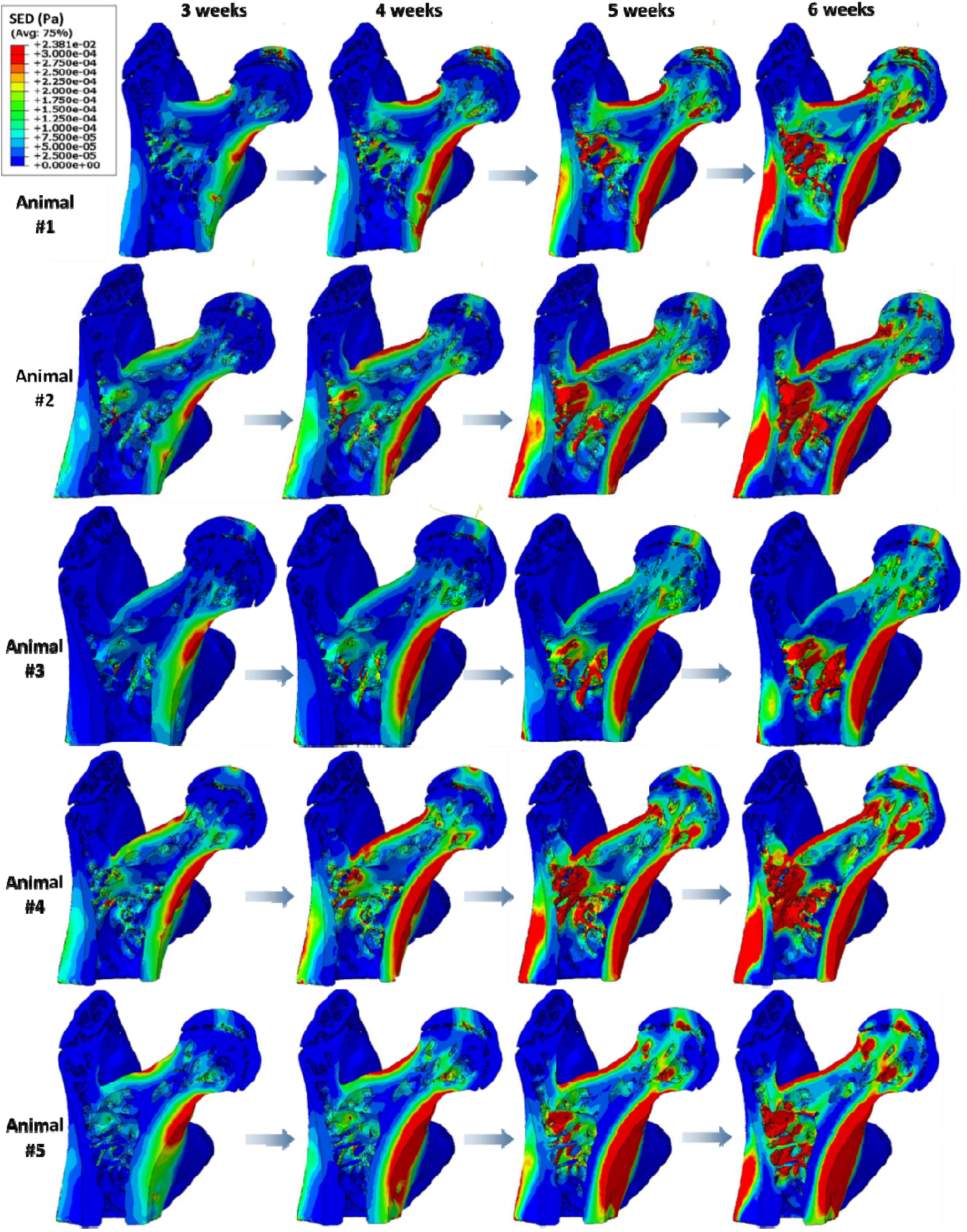
Predicted change in strain energy density distributions from 3 to 6 weeks. Anterior-posterior cross-section views of distributions of SED (Pa) at 3, 4, 5, and 6 weeks of metastasis according to mechanoregulation theory in computational models of proximal femurs.

### 2.6. Statistical Analysis

Statistical analyses were performed using MiniTab (version 17) software. Each parameter was assessed for equal variance (F test) and student t-tests were implemented to determine whether averaged data was statistically significant between groups of equal variances. Welsh’s test was applied where sample groups had unequal variance. Results are displayed as mean ± standard deviation, with significance defined as a p value of < 0.05, and further significance also identified (p<0.01, p<0.001). Error bars in all bar charts represent standard deviation.

## 3. Results

The mechanoregulation algorithm was applied over a three week period from 3 to 6 weeks to predict bone remodelling driven by the evolving mechanical environment within bone tissue during breast cancer-bone metastasis for 5 subjects. We assessed the mechanical environment (strain distribtuion, average maximum strain) (Figure 3), mechanical stimulus (SED) and bone mineral density distribution at the initial timepoint (3 weeks), and at every 7 increments of the algorithm, representing 1 week, to the final time point (6 weeks), see Figure 4 and Figure ***5***.

**Figure 4:**
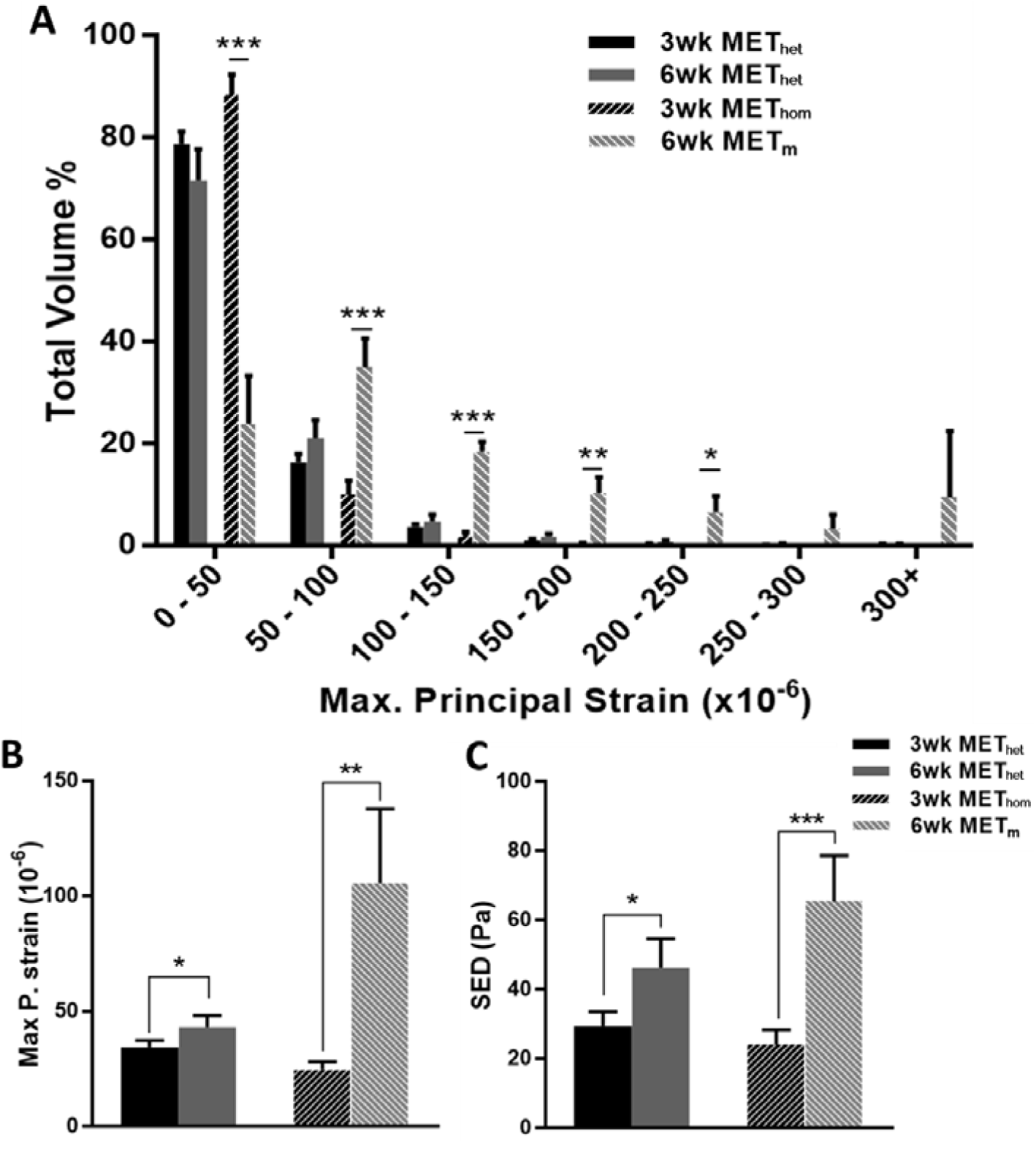
Validation of the initial homogeneous material properties against subject-specific heterogeneous models with resorption rate *C_1_^R^* of 1600. Initially bone tissue was assumed to be an homogeneous material (3wk MET_hom_), which underpredicted the (A) distribution of maximum principal strain as a percentage of bone volume, (B) average maximum principal strain and (C) SED relative to the heterogeneous, micro-CT derived subject-specific models of the same bones at this initial timepoint (3wk MET_het_). **(2) Application of the mechanoregulation theory for 21 increments, representative of 3 weeks of disease development (6wk MET_m_)**. We predicted the evolution of tissue heterogeneity and predicted (A, B) strains and (C) SED were higher than subject-specific heterogeneous models at the same timepoint (6wk MET_het_). *p<0.05, **p<0.01, p***<0.001.

**Figure 5:**
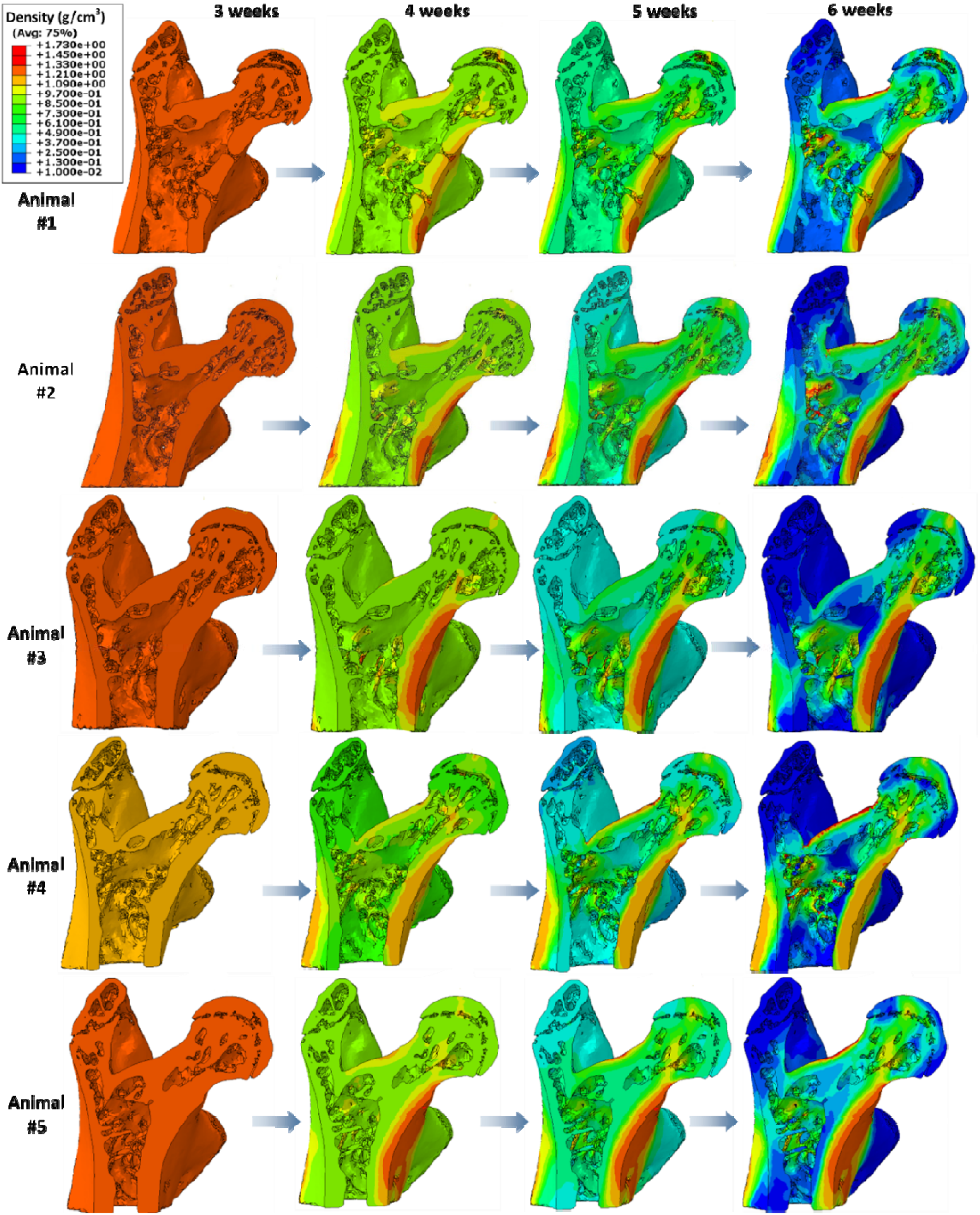
Predicted change in bone mineral density from 3 to 6 weeks. Anterior-posterior cross-section views of bone resorption at 3, 4, 5, and 6 weeks of metastasis according to mechanoregulation theory

### 3.1. Validation of model predictions relative to subject-specific heterogeneous models

The initial homogeneous µCT-FE models of this study (3wk MET_hom_) were validated against micro-CT derived subject-specific models of the same bones at this initial time point (3wk MET_het_). This assumption over-predicted the percentage bone volume at the lowest strains, relative to the heterogeneous models, see Figure 4A. Specifically, the 3 week homogeneous models had a significantly lower average maximum principal strain (3wk MET_hom_: 24.16 ± 3.48% vs. 3wk MET_het_: 34.33 ± 2.9% , p<0.000) and the average mechanical stimulus SED was also significantly lower (3wk MET_hom_: 24.03 ± 2.9% vs. 3wk MET_het_: 29.41 ± 4.1% , p<0.000) see Figure 4B, C. There was an increase in the % bone volume experiencing strains below 50 μe (∼88 %) relative to that predicted by the heterogeneous approach (∼79 %). However, there was a decrease in % bone volume in the 50-100 μe range (∼10% vs. 16%). Thus this assumption represented an initial over prediction (by approximately 3%) of bone tissue volume experiencing strains below the resorption threshold.

Nonetheless, upon application of the mechanoregulation theory the initially homogeneous tissue evolved to become heterogeneous and the mechanical environment evolved in a similar manner to that predicted through heterogeneous subject-specific FE models, as is described in further detail below. The predictive ability of the mechanoregulation theory was also validated against published experimental data, see Supplementary Figure 4.

### 3.2. Evolution of mechanical stimuli within metastatic proximal femurs from 3 to 6 weeks

At week 3 the mechanical stimulus (SED) was highest within the cortical bone tissue of the femoral neck and at the loaded surface of the femoral head for all models (Figure 3). Next, the mechanoregulation algorithm was applied to predict bone remodelling on the basis of a mechanical stimulus (strain energy density) over a period of 21 days, representative of 3 weeks of metastatic disease development. We predicted the evolution in mechanical stimulation (SED) driven by the disruption to the mechanical environment by 3 weeks of tumour metastasis. By week 6 there was in increase in SED within the femoral neck for all subjects.

We analysed the mean maximum principal strain and SED, which were found to increase significantly in both the MET_het_ and MET_hom_ models by 6 weeks (Figure 4B, C). We analysed the strain distributions (6wk MET_m_) and change from week 3 to week 6, and report that there was an increase in strain distribution (i.e. positive skew) as bone tissue is resorbed over this timeframe for both the subject-specific heterogeneous and mechanoregulation models (Figure 4A). The mechanoregulation models predicted a significant decrease in % bone volume in the 0 – 50µ range relative to the initial time point (6wk MET_m_: 23.86 ± 9.36% vs. 3wk MET_hom_: 88.33 ± 3.96%, p<0.001), but a decrease in the % volume in the 50 – 100µ range (6wk MET_m_: 34.88 ± 5.65% vs. 3wk MET_hom_: 9.89 ± 2.8%, p<0.001). Although not significant, for the heterogeneous models (MET_het_) the maximum principal strain distribution decreased in the 0 – 50µ range (6wk MET_het_: 71.63 ± 5.99% vs. 3wk MET_het_: 79.25 ± 2.98%, p = 0.06) and 50 – 100µ range (6wk MET_het_: 20.89 ± 3.68% vs. 3wk MET_het_:15.89 ± 1.98%, p = 0.055).

### 3.3. Bone mineral density within metastatic proximal femurs from 3 to 6 weeks

By 6 weeks application of the bone remodelling algorithm we predicted elevated bone tissue mineral density (calculated from predicted tissue mechanical properties) in these same cortical neck and femoral head regions (Figure 4). It should be noted that, at this 6 week timepoint, the models had not reached homeostasis and remodelling was still active.

By the 6-week time point, application of the remodelling algorithm predicted that 26.51 ± 9.96% of the model volume decreased below a bone mineral density of 0.1 g/cm^3^. In comparison, our previous experimental study reported that bone volume in the trabecular regions of proximal femurs reported from prior micro-CT analysis had decreased by 21.15% between 3 weeks and 6 weeks post-inoculation of metastatic breast cancer cells [6] (see **Error! Reference source not found.**). In addition, the area of lowest density in each model consistently presented in the greater and lesser trochanter regions, as is illustrated in the anterior-posterior cross-sectional views of cortical and trabecular bone (Figure *6*A). This correlates with findings from our experimental study in which osteolysis was reported in the greater trochanter regions by 6 weeks post-inoculation [6] (Figure *6*B, 7C). This region- specific osteolytic destruction of metastatic tissue arose in the exact same areas of the femoral cortical and trabecular space as those reported in our experimental study from which the micro-CT data for these models originated [6], which provides qualitative validation of these computational models. Of note, density was highest in the cortical bone tissues of the femoral neck and the inter-trochanter space (Figure *6*A).

**Figure 6:**
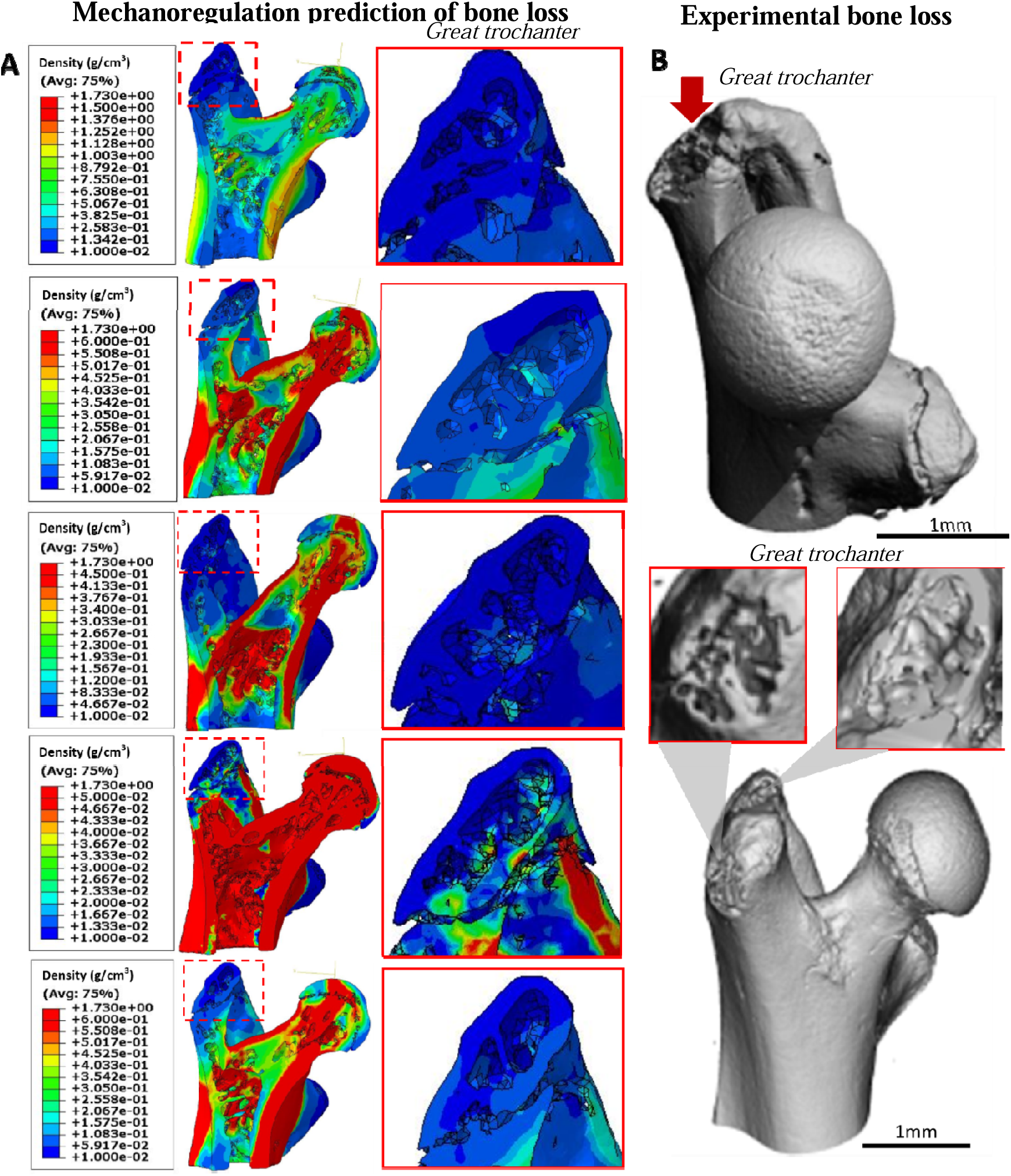
Predicted changes in bone density in the trochanter regions and validation against experimental data. (A) Anterior-posterior cross-section views of predicted bone mineral density for each metastatic model (n=5) after 3 weeks application of mechanoregulation theory (note: the colour legends for density are specific to each model). Insets: Visualisation of predicted resorption in greater trochanter regions. (B) Experimental study reported overt osteolysis in the greater trochanter (in red) by 6 weeks post-inoculation by 3D µCT analysis [6]

### 4.0 Discussion

This study is the first to implement subject-specific finite element models of metastatic bone tissue and apply a computational mechanoregulation framework to investigate whether the development of osteolysis is a function of the altered mechanical environment within bone tissue during breast cancer-bone metastasis. Here we report that, although we implemented a global rate of resorption, we predicted a localised decrease in bone mineral density in the greater trochanter region of the femur, the exact regions where osteolytic lesions were identified in an associated experimental study [6]. Moreover, application of the mechanoregulation algorithm predicted that the spatial mechanical environment evolved in a similar manner to that predicted through subject-specific finite element models over this 3 week time period of metastasis [7]. These results support the hypothesis that early changes in the physical environment of bone tissue during metastasis may elicit mechanobiological cues for bone cells and activate later osteolytic bone destruction.

Some limitations in this study must be considered. Firstly, at the initial stage (3 weeks) bone tissue was assumed to be an isotropic, homogeneous and linear elastic material, which does not fully represent the heterogeneous, anisotropic, non-linear and poroelastic behaviour of bone tissue. Bone tissue is commonly assumed to be linear elastic at low strain, which is relevant to the current study (0 - 300µ , < 0.3%) and below the yield strain (∼3%) of mouse bone tissue. Due to the challenges of implementing mechanoregulation theory in μCT-FE models, linear elasticity and tissue isotropy are often assumed [18, 21, 22], although this assumption might somewhat underestimate the mechanical stimulus (SED) relative to poroelasticity (Falcinelli et al., 2020). We did investigate a combined UMAT and USDFLD subroutine to predict bone remodelling within a heterogeneous model, but this approach proved problematic to implement due to convergence issues arising because of boundary discrepancies between neighbouring elements of differing properties. Thus, we validated our initial (3-week) model predictions against subject-specific micro-CT derived heterogeneous FE models of the same bones [7], and predicted a lower strain distribution and SED for the homogeneous model (see Supplementary Figure 3), which would provide a higher initial stimulus for bone resorption. However, the mechanoregulation theory predicted evolution of the initially homogeneous tissue to become heterogeneous over the simulation period (i.e. by 6 weeks) and, importantly, the mechanical environment evolved in a similar manner to that predicted through heterogeneous subject-specific FE models and although we predicted an increase in bone resorption (∼5%) relative to the experimental study we were able to predict region-specific bone remodelling. Secondly, a parameter variation study was conducted to establish a resorption rate (*C_1_^R^*) that could predict bone loss at a rate consistent with our experimental study, but we used the same rate for cortical and trabecular bone to avoid introducing complex surface interactions not reflective of *in vivo* conditions. Moreover, the models did not account for biochemical signalling between tumour and bone cells, longer- term mechanoregulatory responses or variations in loading angle and magnitude during the murine trotting cycle. Previous studies have successfully coupled biochemical signalling between osteoblasts and osteoclasts with mechanoregulation theory to study bone remodeling [35] or varied resorption and formation rates according to experimental data [21].

Nonetheless, the homogeneous model presented here, in which bone remodelling is governed by mechanical stimuli alone, predicts density and mechanical stimuli evolution comparable to subject-specific finite element models for 6-week-old metastatic animals [6, 7]. Moreover, osteolytic regions predicted by mechanoregulation theory coincided with regions of osteolysis identified by micro-CT in our experimental study [6]. Future studies could investigate the influence of tissue heterogeneity, biochemical signalling and loading on predicted bone tissue resorption, and quantitatively validate the strains by longitudinal Digital Volume Correlation analysis of bone tissue during metastasis [36].

Here we report that the mechanoregulation theory predicted the evolution of the mechanical environment (principal strain, SED) from 3 to 6 weeks, which corresponds with the predictions of micro-CT derived heterogeneous finite element models at 6 weeks [7] and provide further validation of models driven by mechanoregulation theory. This rate implementation predicted that approximately 27% of the model volume would reduce to low bone density, which was within the same range as bone volume fraction reductions in the trabecular space (21.15%) reported experimentally [6]. Bone tissue resorption was more prominent within these models than formation, which is supported by similar studies of female C57BL/6 murine tibiae, wherein bone tissue formation only occurred at high loads (9 - 13N) [31, 37]. In addition, the rate of change of bone density under resorption was higher (*C_1_^R^* = 1600), and formation lower (*C_1_^F^* = 1.79), than that previously determined for healthy tissue (*C_1_^R^* = 15.5, *C_1_* = 45.4, respectively) [14]. It is notable that implementing a lower resorption rate (*C_1_^R^* = 160), predicted an increase in strain distribution, maximum principal strain and SED (Supplementary Fig 3). Further study on the resorption rates could shed light on the evolving mechanical environment and sensitivity to these remodelling parameters.

Physiological loads were applied to subject-specific proximal femur models and a bone mechanoregulation algorithm was implemented to predict element-specific density changes over 21 days. Qualitative analysis revealed that the lowest bone mineral density values were predicted to arise in the greater and lesser trochanter of the femurs. Interestingly, a second cohort of proximal femurs from the same animal model of bone metastasis were experimentally analysed (by micro-CT) 6 weeks after tumour-inoculation and these animals consistently presented with overt osteolytic lesions in the greater trochanter region [6]. Thus, the application of the mechanoregulation theory has accurately predicted the spatial nature of bone resorption within the proximal femur that corresponds to previous experimental results. That bone mineral density was highest along the femoral neck and femoral head, is consistent with the healthy human proximal femur [38] and strain distributions are highest in the femoral neck in a human osteoporotic femur [39]. Interestingly, significant bone tissue resorption was reported by micro-CT analysis of the femoral head regions after 6 weeks of bone metastasis development, and indeed the femoral head was absent in 3 of 7 metastatic proximal femurs [6], possibly due to fracture along the femoral neck. The study implemented a continuum approach to predict bone cell activity in response to mechanobiological cues, but did not explicitly model these cells. Quantification of bone cell and cancer cell populations at different time points throughout metastatic development could validate the predictions. Overall, application of the computational bone mechanoregulation framework presented here demonstrated the spatial nature of resorption by 6 weeks that correlated to the experimental findings.

### 5.0 Conclusion

In conclusion, this study is the first of its kind to predict, using mechanoregulation theory, bone remodelling in µCT-derived finite element models of metastatic bone tissue, prior to the development of overt osteolytic lesions. The bone remodelling algorithm predicted bone mineral density to decrease in regions that coincide with experimental studies of osteolysis. This study also reported changes in strain distribution from 3 weeks to 6 weeks, which were in keeping with predictions of micro-CT derived FE models of femurs at 3 weeks and 6 weeks post-inoculation of metastatic cells. Thus we propose that mechanobiology may play a role in the adaption of the bone tissue extracellular matrix to metastasis and contribute to the later development of osteolysis.

## Competing interests

The authors have no known competing financial interests or personal relationships that could have appeared to influence the work reported in this paper.

## Supplementary Figures

**Supplementary Figure 1:**
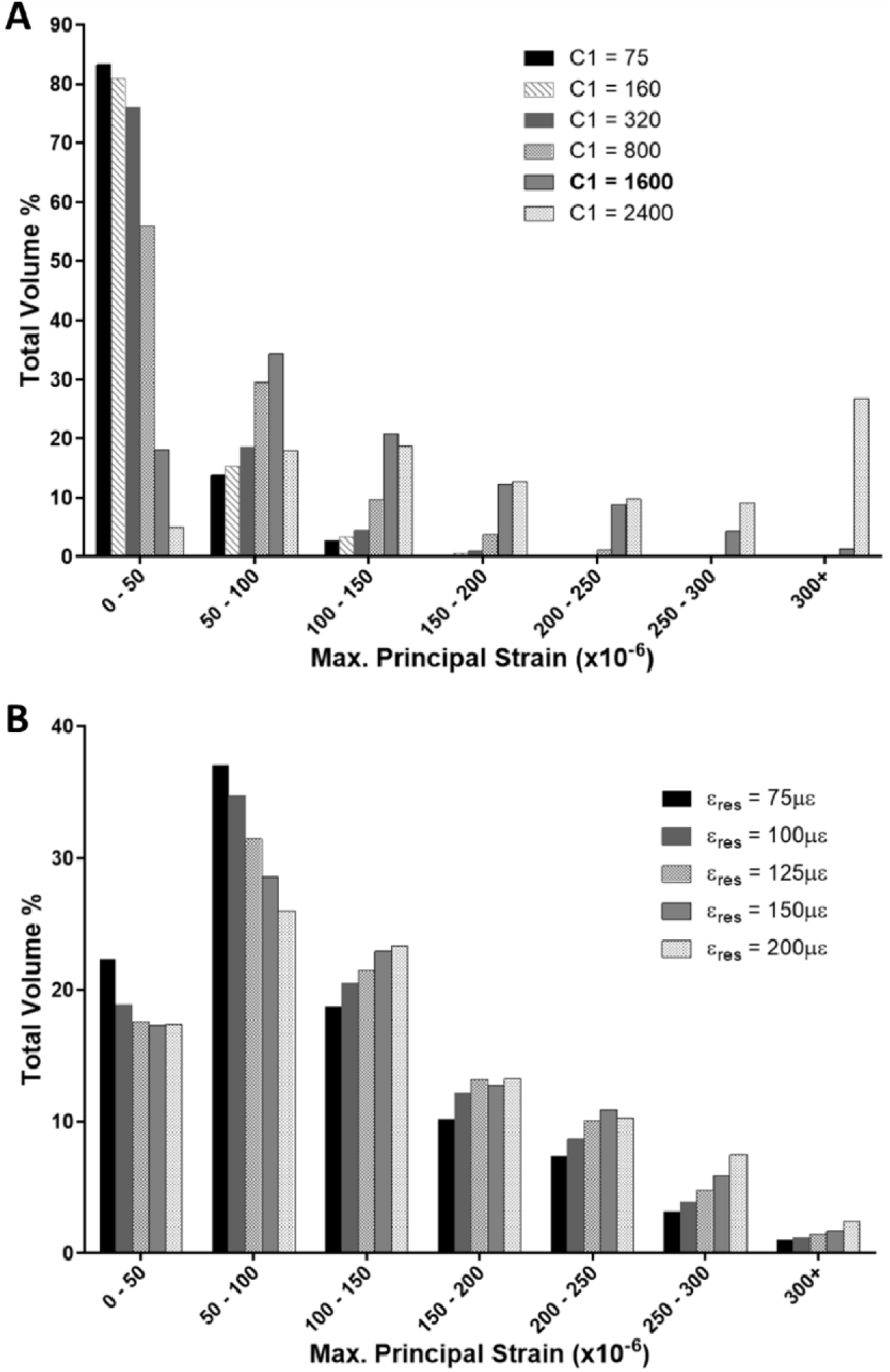
Parameter sensitivity analyses. Distribution of maximum principal strain as a percentage of bone volume in one metastatic proximal femur model, predicted at 6 weeks, comparing between values of (A) bone tissue resorption constant, *C_1_^R^*, and (B) resorption threshold strain value (□_res_) applied to one proximal femur model. The greatest difference in percentage volume for altered □_res_ was between 75 and 200µ□ in the 50-100 range, at 11.09%.

**Supplementary Fig 3:**
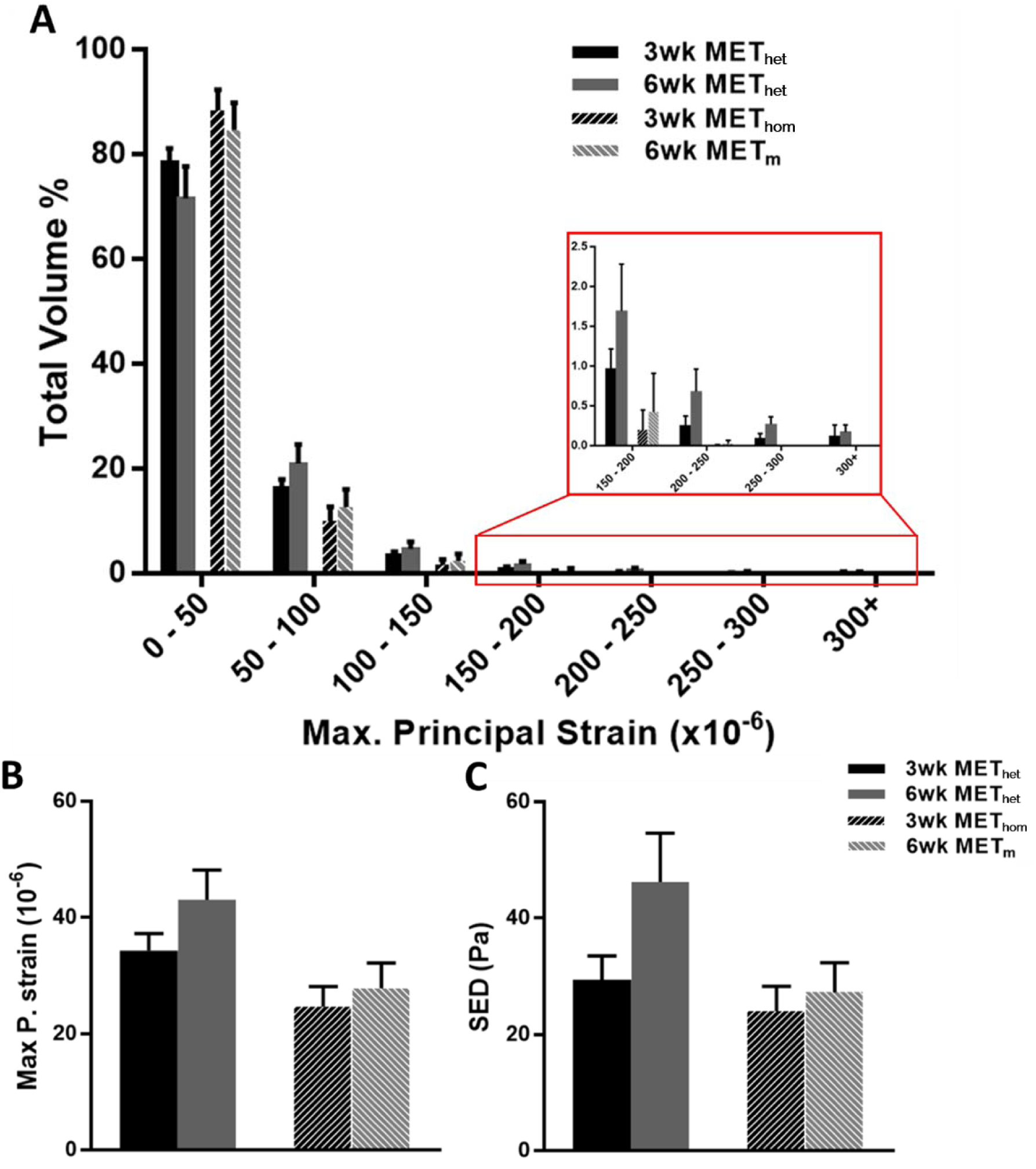
Validation of the initial homogeneous material properties against subject-specific heterogeneous models with a resorption rate C_1_^R^ of 160: **(1)** Initially bone tissue was assumed to be an isotropic, homogeneous and linear elastic material (3wk MET_hom_), which underpredicted the (A) distribution of maximum principal strain as a percentage of bone volume, (B) average maximum principal strain and (C) SED relative to the heterogeneous, micro-CT derived subject-specific models of the same bones at this initial timepoint (3wk MET_het_). **(2)** Following application of the mechanoregulation theory for 21 increments, representative of 3 weeks of disease development (6wk MET_m_), we predicted the evolution of tissue heterogeneity and predicted (A, B) strains and (C) SED were lower than subject-specific heterogeneous models at the same timepoint (6wk MET_het_).

**Supplementary Figure 4:**
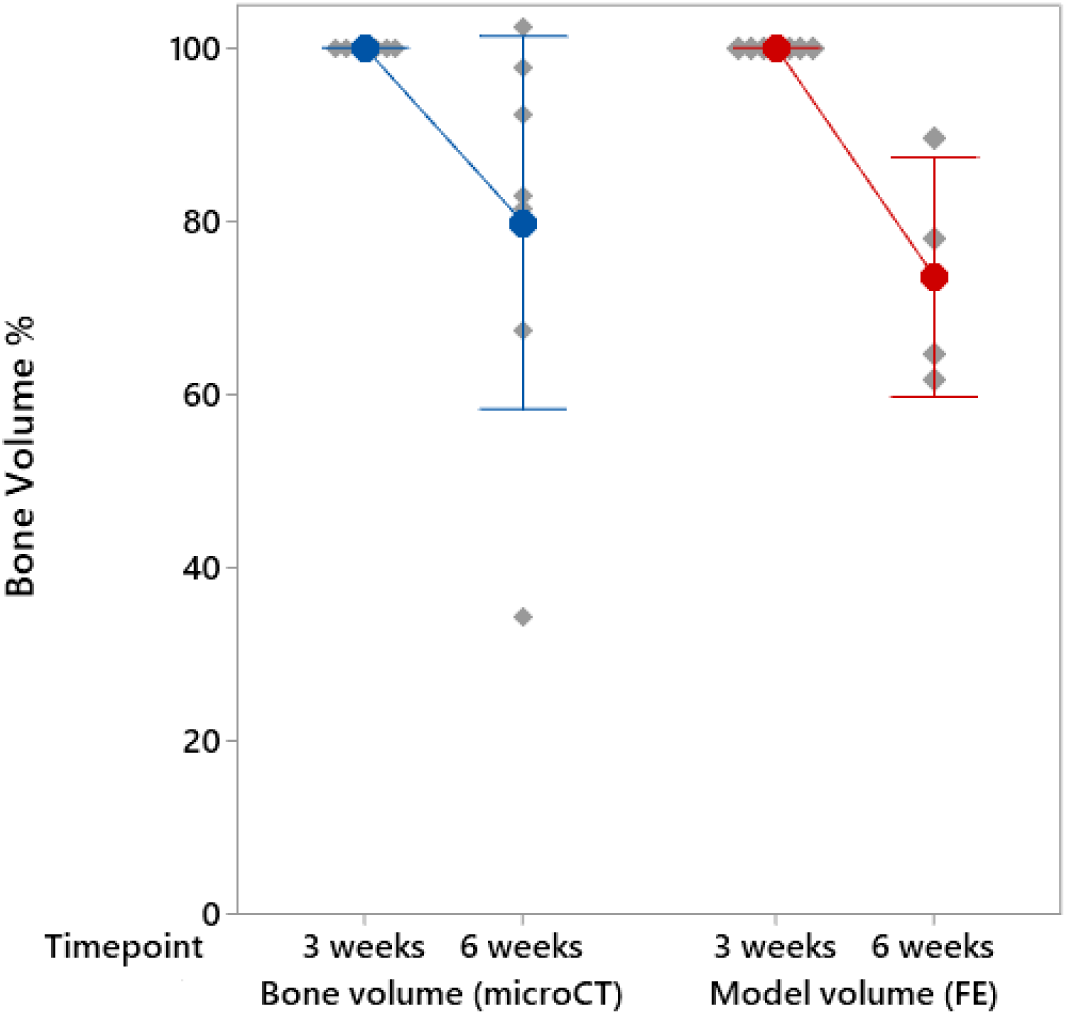
Validation of computational predictions of bone resorption based on mechanoregulation theory to experimental results. The change in bone volume (BV) from week 3 to week 6 was measured experimentally for metastatic femora (in blue) by micro-CT and was found to ∼ 21% [6], whereas the mechanoregulation theory predicted the decrease in bone volume (i.e. elements above bone density) after 21 iterations (6 weeks) to be 26.51 ± 9.96%. Interval plots present mean, standard deviation and datapoints from each individual femur. Note: while furthest from the rest, the lowest datapoint (34.09%) was not a significant outlier from its group according to Grubb’s test.

**Supplementary Fig 7:**
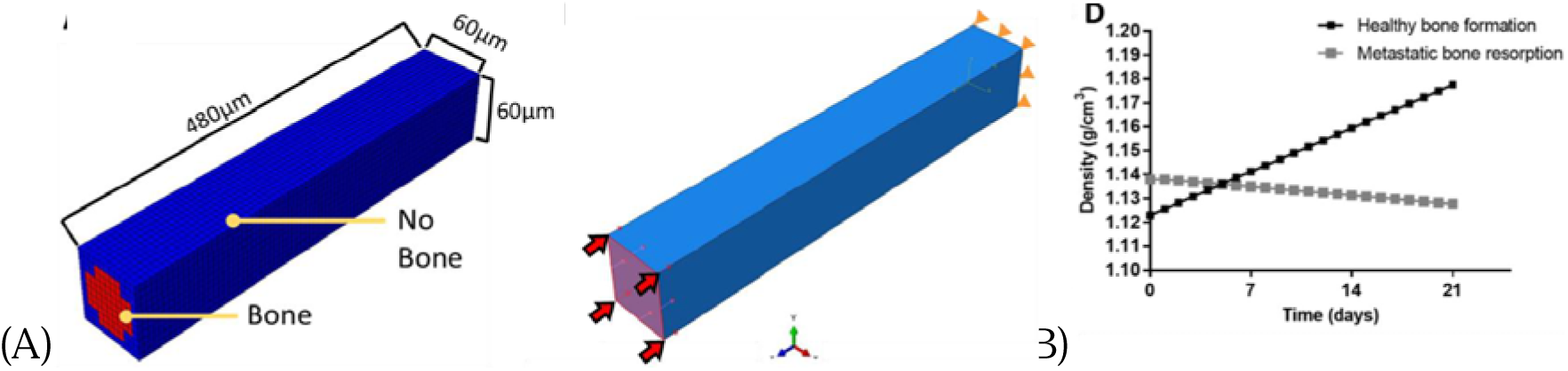
Derivation of remodelling rates (C_1_). The mechanoregulation theory was applied to (A) a simplified model of an individual metastatic trabecula under physiological loading. (B) The average predicted change in bone mineral density (M_mean_) over 3 weeks (21 increments) was representative of experimental changes over the same timeframe [6]. Specifically (1) the time constant for bone formation (C_1_^F^) was dervied at high strain (1250μ□) to predict an overall density increase (1.123 g/cm^3^ to 1.178 g/cm^3^) from healthy bone and (2) the bone resorption constant (C_1_^R^) was derived such that an overall density decrease (1.138 g/cm^3^ to 1.128 g/cm^3^) arose, capturing experimental rates of osteolysis in metastatic bone [6].

## Notes

### Competing Interest Statement

The authors have declared no competing interest.

## References

[1] S. Paget, The distribution of secondary growths in cancer of the breast, Lancet (1889) 571–573.

[2] T.A. Guise, The vicious cycle of bone metastases, J Musculoskelet Neuronal Interact 2(6) (2002) 570–2.

[3] S.A. Arrington, J.E. Schoonmaker, T.A. Damron, K.A. Mann, M.J. Allen, Temporal changes in bone mass and mechanical properties in a murine model of tumor osteolysis, Bone 38(3) (2006) 359–67.

[4] A. Nazarian, D. von Stechow, D. Zurakowski, R. Muller, B.D. Snyder, Bone volume fraction explains the variation in strength and stiffness of cancellous bone affected by metastatic cancer and osteoporosis, Calcif Tissue Int 83(6) (2008) 368–79.

[5] L. Richert, L. Keller, Q. Wagner, F. Bornert, C. Gros, S. Bahi, F. Clauss, W. Bacon, P. Clézardin, N. Benkirane-Jessel, F. Fioretti, Nanoscale Stiffness Distribution in Bone Metastasis, World Journal of Nano Science and Engineering 05(04) (2015) 219–228.

[6] A.S. Verbruggen, E.C. McCarthy, R.M. Dwyer, L.M. McNamara, Temporal and spatial changes in bone mineral content and mechanical properties during breast-cancer bone metastases, Bone Reports (2022) 101597.

[7] A.S. Verbruggen, L.M. McNamara, Mechanoregulation may drive osteolysis during bone metastasis: A finite element analysis of the mechanical environment within bone tissue during bone metastasis and osteolytic resorption, Journal of the Mechanical Behavior of Biomedical Materials (2023) 105662.

[8] Y. Kim, H.G. Othmer, Hybrid models of cell and tissue dynamics in tumor growth, Math Biosci Eng 12(6) (2015) 1141–56.

[9] K.A. Rejniak, A.R. Anderson, Hybrid models of tumor growth, Wiley Interdiscip Rev Syst Biol Med 3(1) (2011) 115–25.

[10] X. Zhou, J. Liu, A computational model to predict bone metastasis in breast cancer by integrating the dysregulated pathways, BMC Cancer 14 (2014) 618.

[11] L.M. Cook, A. Araujo, J.M. Pow-Sang, M.M. Budzevich, D. Basanta, C.C. Lynch, Predictive computational modeling to define effective treatment strategies for bone metastatic prostate cancer, Sci Rep 6 (2016) 29384.

[12] A. Araujo, L.M. Cook, C.C. Lynch, D. Basanta, An integrated computational model of the bone microenvironment in bone-metastatic prostate cancer, Cancer Res 74(9) (2014) 2391–401.

[13] P. Tracqui, Biophysical models of tumour growth, Reports on Progress in Physics 72(5) (2009).

[14] L.M. McNamara, P.J. Prendergast, Bone remodelling algorithms incorporating both strain and microdamage stimuli, Journal of biomechanics 40(6) (2007) 1381–1391.

[15] B.M. Mulvihill, L.M. McNamara, P.J. Prendergast, Loss of trabeculae by mechano- biological means may explain rapid bone loss in osteoporosis, Journal of The Royal Society Interface 5(27) (2008) 1243–1253.

[16] R. Huiskes, H. Weinans, H. Grootenboer, M. Dalstra, B. Fudala, T. Slooff, Adaptive bone-remodeling theory applied to prosthetic-design analysis, Journal of biomechanics 20(11-12) (1987) 1135–1150.

[17] F.A. Schulte, F.M. Lambers, D.J. Webster, G. Kuhn, R. Müller, In vivo validation of a computational bone adaptation model using open-loop control and time-lapsed micro-computed tomography, Bone 49(6) (2011) 1166-1172.

[18] F.A. Schulte, A. Zwahlen, F.M. Lambers, G. Kuhn, D. Ruffoni, D. Betts, D.J. Webster, R. Müller, Strain-adaptive in silico modeling of bone adaptation—a computer simulation validated by in vivo micro-computed tomography data, Bone 52(1) (2013) 485–492.

[19] A.F. Pereira, B. Javaheri, A. Pitsillides, S. Shefelbine, Predicting cortical bone adaptation to axial loading in the mouse tibia, Journal of the Royal Society Interface 12(110) (2015) 20150590.

[20] C. Villette, J. Zhang, A. Phillips, Influence of femoral external shape on internal architecture and fracture risk, Biomechanics and Modeling in Mechanobiology 19(4) (2020) 1251–1261.

[21] V.S. Cheong, A. Campos Marin, D. Lacroix, E. Dall’Ara, A novel algorithm to predict bone changes in the mouse tibia properties under physiological conditions, Biomechanics and Modeling in Mechanobiology 19(3) (2020) 985–1001.

[22] V.S. Cheong, B.C. Roberts, V. Kadirkamanathan, E. Dall’Ara, Bone remodelling in the mouse tibia is spatio-temporally modulated by oestrogen deficiency and external mechanical loading: A combined in vivo/in silico study, Acta Biomaterialia 116 (2020) 302–317.

[23] C. Quinn, A. Kopp, T.J. Vaughan, A coupled computational framework for bone fracture healing and long term remodelling: Investigating the role of internal fixation on bone fractures, Interna^LJ^tional Journal for Numerical Methods in Biomedical Engineering 38(7) (2022) e3609.

[24] A. Sohail, M. Younas, Y. Bhatti, Z. Li, S. Tunç, M. Abid, Analysis of trabecular bone mechanics using machine learning, Evolutionary Bioinformatics 15 (2019) 1176934318825084.

[25] A. Ramos, J. Simoes, Tetrahedral versus hexahedral finite elements in numerical modelling of the proximal femur, Medical Engineering & Physics 28(9) (2006) 916–924.

[26] J.P. Charles, O. Cappellari, J.R. Hutchinson, A dynamic simulation of musculoskeletal function in the mouse hindlimb during trotting locomotion, Frontiers in Bioengineering Biotechnology 6 (2018) 61.

[27] Y. Lu, D. Zuo, J. Li, Y. He, Stochastic analysis of a heterogeneous micro-finite element model of a mouse tibia, Journal of Medical Engineering and Physics 63 (2019) 50–56.

[28] H.M. Frost, Bone “mass” and the “mechanostat”: a proposal, The anatomical record 219(1) (1987) 1–9.

[29] P.T. Scannell, P.J. Prendergast, Cortical and interfacial bone changes around a non- cemented hip implant: simulations using a combined strain/damage remodelling algorithm, Medical Engineering & Physics 31(4) (2009) 477–488.

[30] B. Van Rietbergen, R. Huiskes, H. Weinans, D. Sumner, T. Turner, J. Galante, The mechanism of bone remodeling and resorption around press-fitted THA stems, Journal of biomechanics 26(4-5) (1993) 369–382.

[31] R. De Souza, M. Matsuura, F. Eckstein, S. Rawlinson, L. Lanyon, A. Pitsillides, Non-invasive axial loading of mouse tibiae increases cortical bone formation and modifies trabecular organization: a new model to study cortical and cancellous compartments in a single loaded element, Bone 37(6) (2005) 810–818.

[32] C.H. Turner, M. Forwood, J.Y. Rho, T. Yoshikawa, Mechanical loading thresholds for lamellar and woven bone formation, Journal of Bone and Mineral Research 9(1) (1994) 87–97.

[33] H. Razi, A.I. Birkhold, R. Weinkamer, G.N. Duda, B.M. Willie, S. Checa, Aging leads to a dysregulation in mechanically driven bone formation and resorption, Journal of Bone and Mineral Research 30(10) (2015) 1864–1873.

[34] D.M. Geraldes, A.T. Phillips, A comparative study of orthotropic and isotropic bone adaptation in the femur, International journal for numerical methods in biomedical engineering 30(9) (2014) 873–889.

[35] R. Hambli, Connecting mechanics and bone cell activities in the bone remodeling process: an integrated finite element modeling, Frontiers in Bioengineering and Biotechnology 2 (2014) 6.

[36] M. Palanca, S. Oliviero, E. Dall’Ara, MicroFE models of porcine vertebrae with induced bone focal lesions: Validation of predicted displacements with digital volume correlation, J Mech Behav Biomed Mater 125 (2022) 104872.

[37] A.M. Weatherholt, R.K. Fuchs, S.J. Warden, Cortical and trabecular bone adaptation to incremental load magnitudes using the mouse tibial axial compression loading model, Bone 52(1) (2013) 372–379.

[38] E. Dall’Ara, R. Eastell, M. Viceconti, D. Pahr, L. Yang, Experimental validation of DXA-based finite element models for prediction of femoral strength, Journal of the mechanical behavior of biomedical materials 63 (2016) 17–25.

[39] E. Verhulp, B. van Rietbergen, R. Huiskes, Load distribution in the healthy and osteoporotic human proximal femur during a fall to the side, Bone 42(1) (2008) 30–5.

